# Observing picomolar protein unfolding using resonance light scattering

**DOI:** 10.1101/2024.11.20.624557

**Authors:** Alain Bolaño Alvarez, Kristian B. Arvesen, Kasper F. Hjuler, Peter Bjerring, Steffen B. Petersen

## Abstract

We here present a novel and sensitive methodology for determining the melting point (MP) of Bovine Serum Albumin (BSA) from micromolar to picomolar concentration levels under label free conditions. At 1 pM we could model the melting with a sharp gaussian. However, from the transient state observed during the melting process by using a simple exponential decay model we determined a time constant of 67 seconds. We applied this methodology by studying a 3.3 pM sample of a botulinum toxin A (BoNT-A) (stabilized with 2.8 nanomolar denatured Human Serum Albumin (HSA)). We were able to determine the Tm of BoNT-A in the presence of the approximately 1000-fold more concentrated HSA.

**Entry for the Table of Contents:** 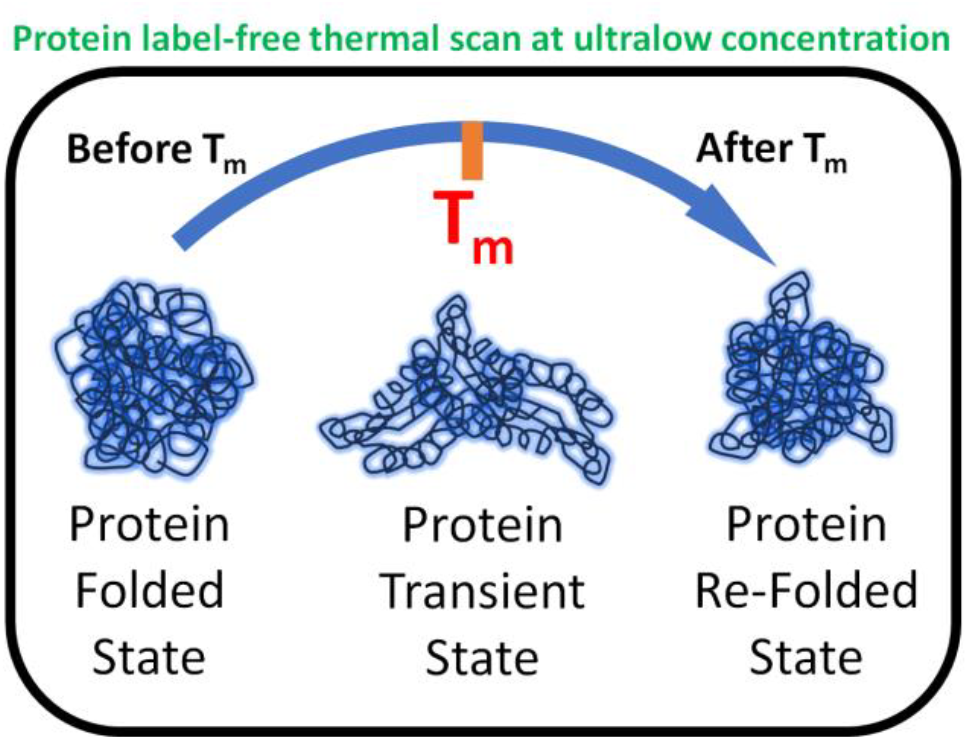

Protein label-free melting point (MP) determination at ultralow concentrations is a huge problem which concern to the biopharmaceutical industry. Here, we present a novel method to determine the MP of bovine serum albumin (BSA) from 1μM to 1pM under label-free conditions. The benefits of this study match the purposes of stability studies in formulations, in which the protein active component is successful at very low concentrations, such as botulinum toxin A (BoNT-A). We used BOCOUTRE, a commercially available pharmaceutical product based on BoNT-A, and Human Serum Albumin (HSA) as a stabilizer. Our method can detect the MP of the stabilizer protein, even if its concentration is markedly different from that of the active component protein (1000-fold) in the case of BOCOUTRE.

---

The detection of protein unfolding at ultralow concentrations is crucial in biopharmaceutical studies. Previous investigations have used fluorescent dyes to enhance the signal-to-noise ratio to achieve detection in the millimolar (mM) range ^[1]^ and nanomolar range ^[2]^. But the dye used could potentially modulate both the transition and stability of the protein as well ^[3]^. We are dealing with a special case of scattering that is called Resonance Light Scattering (RLS) to study protein folding dye-free. This phenomenon is observed if the excitation wavelength used is the same or near the wavelength of absorption of the chromophore and particularly the magnitude of effect increases in aggregates states ^[4]^ which can enhance the observed change in response to the protein melting process. In the present study we have established unequivocal evidence that RLS can be used to monitor the ‘melting’ of selected proteins down to micromolar concentrations using dye-free approach. The methodology monitors the experimental melting temperature as a function of protein concentrations form 1μM to 1pM, under thermal scan range from 20°C to 75°C (optimal optical properties of the cuvettes) and/or 20°C to 90°C, (**Fig. 2, S2 and S3**). In structural studies of proteins, detection at ultralow concentrations is challenging. Our results indicate effective detection showing a clear tendency to decreasing signal-to-noise ratio as the BSA concentration decreases being more than unity ^[5]^ (**Table 1 and Supporting Information**).

**Table 1:** S/N ratio of BSA concentrations. Fluorescence emission at 20 °C (λ_exc_ = 295nm)

| [BSA] | S/N ratio | Intensity at 350 nm |
| --- | --- | --- |
| 1 μM | 1.37 | 411192.7 |
| 1 nM | 1.19 | 10821.6 |
| 1 pM | 1.02 | 5186.3 |

**Figure 1:**
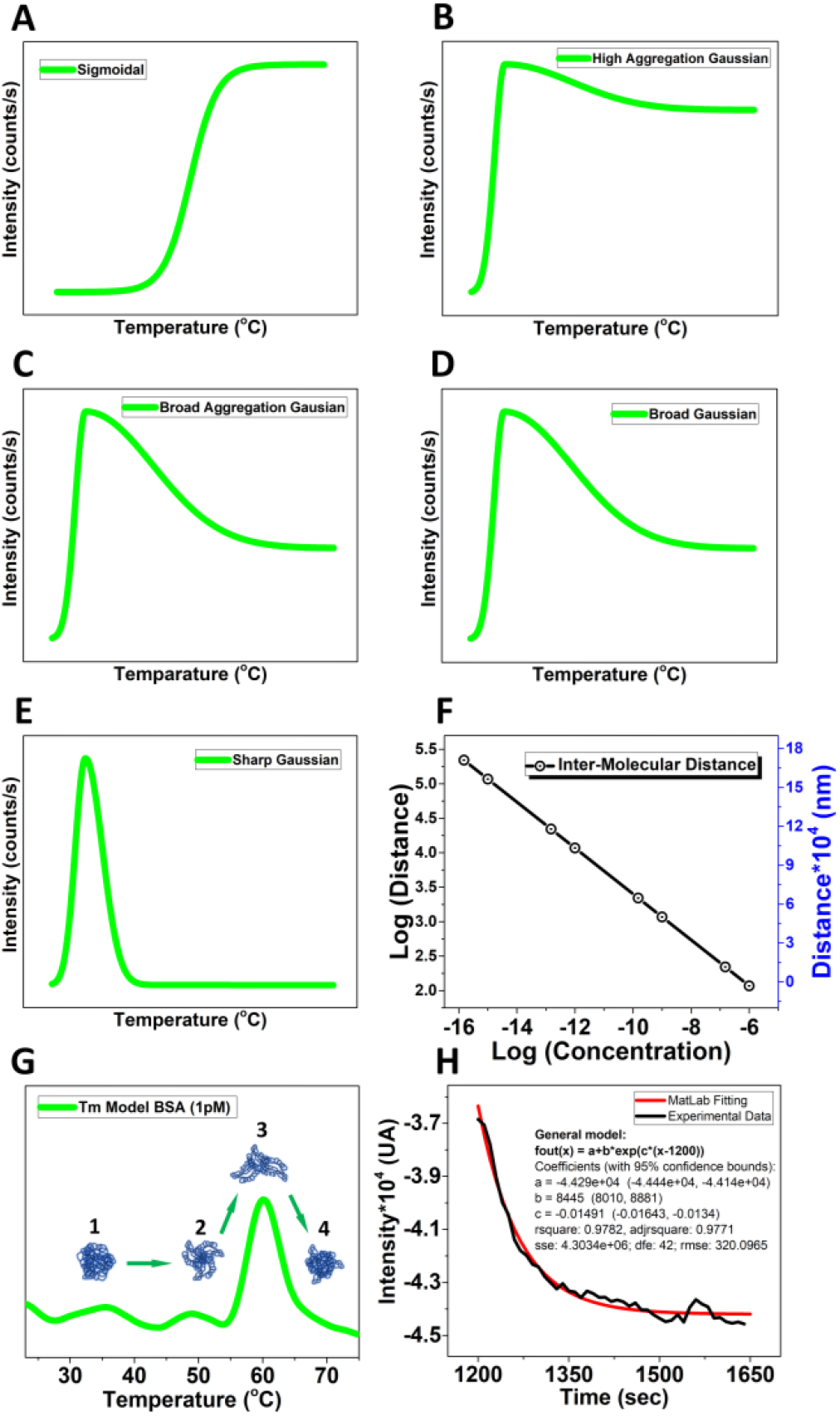
Mathematical Model from MATLAB. The mathematical model contemplates the concentration values fleeting from a Sigmoidal (A), High aggregation gaussian (B), Broad aggregation gaussian (C), Broad gaussian (D), and Sharp gaussian (E). Molecular distance change respect to the concentration (F), Proposed model to explain the experimental observed behavior at 1pM. (1) native BSA structure, (2) before unfolded BSA structures, (3) unfolded BSA structures, (4) refolded BSA structure (new globular structure) (G), Exponential decay kinetic describing the collapsing protein structure into the new globular conformation after unfolding at Tm (see General model parameters in the insert) (H).

**Figure 2:**
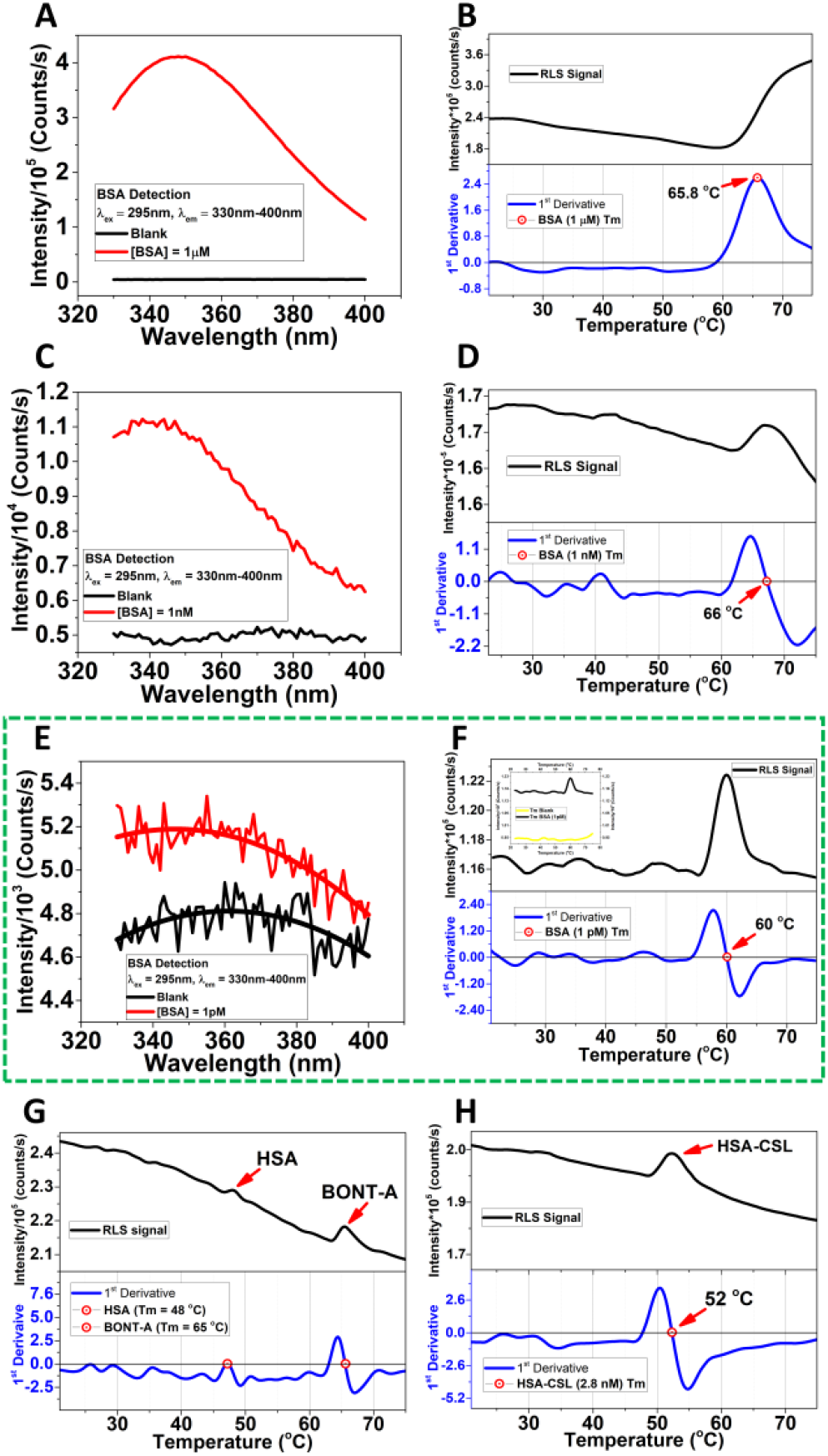
Melting point determination and detection of BSA at ultralow concentration without fluorescent dyes. BSA detection (A) and MP (B) at 1μM, BSA detection (C) and MP (D) at 1nM (69 ± 3°C), and BSA detection (E) and MP (F) at 1pM concentrations, the insert remarks the difference between blank and 1pM BSA concentartion. The excitation wavelength was at 295 nm and emission range was between 330 nm to 400 nm. The average melting point curve corresponding to BSA at 1pM (59.09 ± 1.28°C, After Correction: 61.79 °C) the insert shows the signal respect to the blank (F). Melting point of the BoNT-A (65°C) at 3.3 pM and HSA (48°C) at 2.8 nM in BOCOUTURE (G). Melting point of the HSA (52°C) at 2.8 nM of concentration determined by the standard calibration curve by using a commercial HSA-CSL sample (H). The thermal scans were in the optimal optical properties of the cuvettes. See SI for scans in the range 20 – 90 °C.

When a folded protein is experiencing a progressive heating from close to 0 °C to nearly 100 °C it most likely will ‘melt’ at some intermediate temperature. This melting can be perceived as a phase transition, the proteins folding state changes from the initial state to a new state within a few degrees Celsius. Most often this is an irreversible process, cooling does not regain the original low temperature folded state. Somehow the protein is stuck in new constraints which cannot overcome in the unfolded state where now populates. Various spectroscopic methods can monitor this process: circular dichroism, fluorescence spectroscopy, infrared spectroscopy, and light scattering. Traditionally, determination of the melting point of proteins such as Bovine Serum Albumin (BSA) has been confined to relatively high concentrations, where the cooperative unfolding transition is readily detectable ^[6]^, which effect we observed clearly at 1μM (**Fig. 2B**). Here, we determined the melting point of Bovine Serum Albumin (BSA) at different concentration values from 1pM up to 1μM by RLS, exciting at 295 nm and scanning from 20 – 75 °C and/or 20 – 90 °C (**see Supporting Information**), the **Tm** was corrected by plotting the data from **Table 2**: **Correction = (Tm*1.05)-0.25**, where **Tm** is the experimental melting point in °C, **1.05** is the slope of the plotted data, and **-0.25** is the intercept of the plotted data. This correction improves the accuracy of the melting point values reducing the error introduced by the instrument temperature control system in the holder. The accuracy of the set temperature in the holder was checked using an electronic temperature tester.

**Table 2:** Accuracy of the temperature set at the Peltier temperature controller

| Instruments | Temperature Range (°C) |  |  |  |  |  |  |  |
| --- | --- | --- | --- | --- | --- | --- | --- | --- |
| PT100 | 5 | 15 | 25 | 35 | 45 | 55 | 65 | 75 |
| Peltier | 5 | 15 | 24 | 33 | 43 | 52 | 62 | 72 |

Certainly, at 1pM the self-aggregation is likely to be less pronounced (**Fig. 2F**). However, most protein contain multiple aromatic residues that sense the excitation state of the surrounding aromatic residues ^[7]^ which we claim is relevant to detect the MP at ultralow concentration. We may enhance the scattering intensity with a factor of up to 10-fold because of the many tryptophan will contribute to the observed resonance scattering effect by mimicking a self-aggregation state due to their proximity in the same protein structure (**Scheme 1**). Our methodology provides a visual representation of the unfolding/refolding transitions of BSA molecules, proposing a model to explain this behavior at ultralow concentrations (**Fig. 1E, 1G and 1H**).

We can support our experimental findings by comparing the experimentally determined melting points with predictions obtained using mathematical tools (**Fig. 1**). To explain the proposed model, our data provide distinctive changes in the RLS profile when the temperature increases. These alterations in the light scattering intensity correspond to the structural transitions within the BSA molecules, determining different levels of self-exposition of **Trp**, especially at the temperature range over which the melting occurs, which is centered around 60 °C (**Fig. 2F**). Interestingly we are also able to evaluate the time needed for the transient unfolding event (**τ**_**1pM**_ **= 67**,**07 sec and τ**_**1nM**_ **= 529 sec**), since we can time the duration of the enhanced resonance scattering signal by using a single exponential decay (**see Supporting Information**). We have measured this at all concentrations and the result is shown in **Fig. S1**.

According to our understanding, no other label free method for studying protein melting has been reported for concentrations as low as picomolar concentrations.

**Scheme 1:**
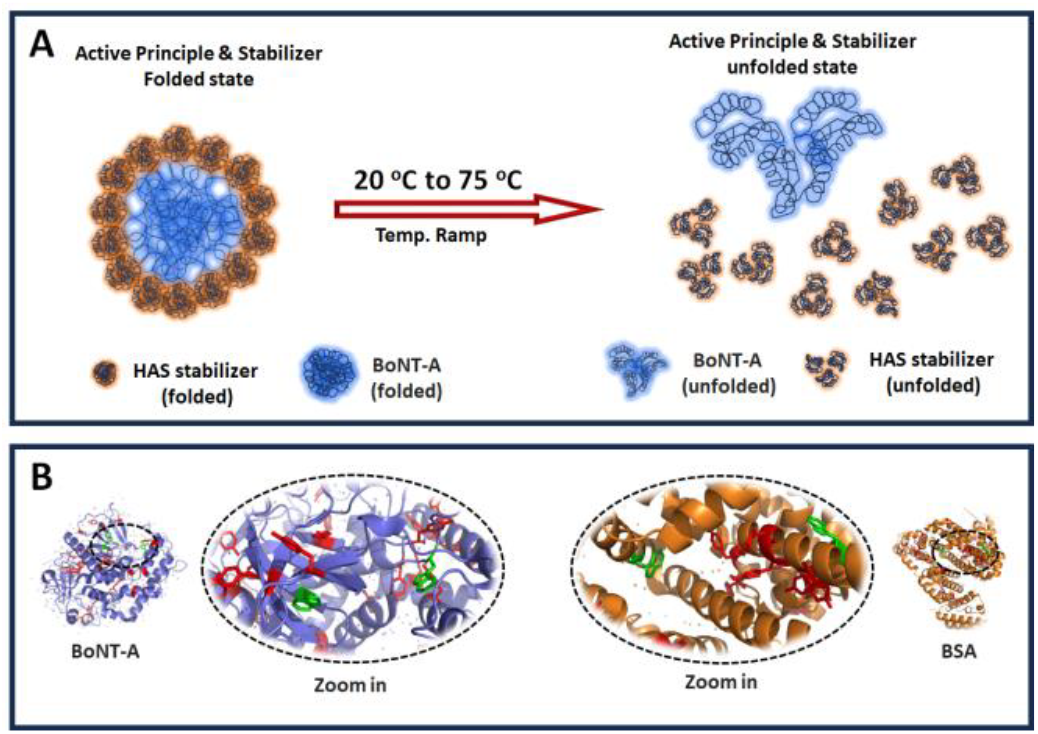
Temperature effect on the protein conformation and relative chromophores proximity. A: Schematic representation of the temperature effect on the active principle & stabilizer folded structure. (See Supporting Information for stabilizer effect). B: Relative proximity between Tyrosine (red) and Tryptophan (green) amino acids. The average distance in between Trp residues is 49.35 Å (BSA, PDB: 4f5s). Pymol was used for visualization.

Currently this method is therefore unique – and we postulate that it may be the only known method where the protein melting process can be studied without aggregational processes contributing to the thermodynamics of the melting process at extremely dilution. In other words – the apparent melting point is most likely shifted because of the aggregation stabilizing the unfolded state at high concentration, which relates to available studies ^[8]^ We have monitored the resonance light scattering method at a range of concentrations. At low concentrations, the light scattering trace appear as a skewed gaussian with a sharp onset and a slow trailing decay. The sharp onset we believe is caused by sudden increase in the physical size of the protein because of a partial unfolding. The trailing decay we suggest is the melted protein seeking a new possibly globular but misfolded conformation (**Fig. 1G and 2F**).

When concentration increases, the trailing decay is observed to increase in time – indicating a more complex search for a new conformation. Increasing the concentration further, we start observing that the protein retains its higher scattering response – indicating that its melted, deformed size does not relax back to the levels observed at lower protein concentration.

However, the time constant cannot be determined at high concentration (1μM) since aggregation process dominance over refolding (**Fig. 2**). Although other methodologies that involved fluorescent dyes can detect proteins in the fM range ^[9]^ they cannot provide information about the conformational state of the protein as we can do with our free dye approach.

In this study to demonstrate our melting point determination dye-free approach, we used a Botulinum Toxin (3.3pM) formulation (BOCOUTURE) which contain HSA (2.8 nM) as a stabilizer (**Supporting Information**). The HSA concentration in BOCOUTURE was determined by constructing a standard calibration curve employing a commercially available HSA-CLS (Behring GmbH) (**Supporting Information**). The analysis revealed two distinct melting points within the BOCOUTURE formulation. The first was observed at 48 °C and the second at 65 °C (**Fig. 2G**). To identify the melting point of HSA within the BOCOUTURE, a sample of HSA-CLS was subjected to RLS under the same experimental conditions (**Supporting Information**). The result showed that the melting point of HSA-CLS was 52 °C (**Fig. 2H**). We have corrected the melting point values by using our previous proposed model (**Table 2 and 3**).

**Table 3:** Melting point after correction.

| Sample | HSA CSL | HSA | BoNT-A |
| --- | --- | --- | --- |
| Exp. melting point ($^\circ\text{C}$) | 52 | 48 | 65 |
| Correction ($^\circ\text{C}$) | 54.05 | 49.89 | 67.57 |

The change in the RLS signal to determine the melting points of the proteins matches the proposed mathematical model, supporting those proteins undergoes a globular conformational re-arrangement that differs from the native structure, which explains the recovered scattering intensity at range 1nM and 1pM (**Fig. 2D, F, G, H**). Our analysis at 1pM did not include aggregation because the intermolecular distance at 1pM (117274.3 nm) makes aggregation extremely unlikely (**Fig. 1A**). We assume that the unfolding is truly a mono-molecular process resulting in a transient change in the protein size, followed by a relaxation to a new non-functional folding state, even if the stabilizer (HSA) forms a shell to protect the active protein (BoNT-A) of self-aggregation, disassembling such scaffold by the temperature scan, thus detecting separately both melting points (**Scheme 1A and Fig. 2G**).

On the other hand, the combination of vacuum degassing and preheating of the buffer solution effectively eliminated bubbles and gas-related interferences, significantly enhancing the signal quality in melting point determination by RLS analyses.

These improvements make these methods valuable for spectroscopic applications to detect molecules at ultralow concentrations.

We believe that our result is important of mainly 2 reasons: 1-we are observing the unfolding event without any dye being introduced into the sample. 2-we can observe the label free unfolding event at a concentration level not achieved by any other biophysical method. Interestingly we are also able to evaluate the time needed for the transient unfolding event, since we can time the duration of the enhanced resonance scattering signal. We have measured this at all concentrations and the result is shown in **Fig. 2**. This highlights how ultralow concentration affects the stability of proteins, such as BSA, HAS and BONT-A. This methodology is innovative and useful to study biopharmaceuticals formulations where BSA or HSA are used as a stabilizer ^[10]^ as well as formulations based on proteins that are extremely diluted, such as botulinum toxin ^[11]^. However, if the stabilizer is a molecule with low molecular weight such as tryptophanate (tryptophan anion) and caprylate (octanoic acid) as occurs in the case of HSA-CSL preparation ^[12]^ some differences in the melting point can emerge. It was the reason for the observed the differences between the melting points of HSA-CLS (52 °C) and HSA stabilizer within the BOCOUTURE formulation (48 °C) (**Fig. 2G-H and Table 1**). However, the conformational behavior of both proteins (BONT-A and HSA) from the folded state to a refolded state remain differentiable and which are well detected by this methodology. Our study can help to understand how stabilizing molecules impact the melting point to trigger new protein conformational behavior (refolding) during thermal stability analysis.

## Signal intensity changes in the focal volume

Modern photon sensing technology has progressed in the most impressive manner. Many cameras declare a quantum efficiency of 0.5 or better for some range of wavelength ^[13]^. When considering the full spectral width, the quantum efficiency may drop to 0.1 at the least sensitive wavelength ^[14]^. In simple absorption terms, we can detect single molecules, if we consider fluorescence the exciting photons is covered as part of the fluorescence process to a longer wavelength photon. Most often the wavelength shift is relatively small, thus the quantum efficiency is nearly unchanged. If we consider the folding/unfolding process of proteins, we will excite the molecule periodically and monitor the absorption or fluorescence as a function of time. Here we expect a near exponential growth or disappearance of a signal whose intensity assuming is indicative of the population of the molecule in a particular folding state. This change in signal intensity must stem from photon detection from a statistical subset of molecules (**Figure 2 and S1**). If we assume that the focal volume we are able to detect a signal from is 3000uL, and the concentration of our molecule of interest is 1pM, we are observing the signal from N = 1.8*10^^9^ molecules.

## Conclusion

This novel approach avoids interferences introduced by fluorescent dye and offers a novel insight for ultrasensitive protein analysis in the pharmaceutical industry, especially at ultralow protein concentrations highlighting that the globular structure of the protein persists even passing the melting point. The reduction in bubble content resulted in improved scattering measurements with improved signal-to-noise ratios. This methodology is sensitive to detecting thermally induced conformational changes, even when the concentration and molecular weight of the stabilizer protein differs dramatically of the active pharmaceutical protein. Thus, it is a versatile technique for detecting both aggregation and conformational changes (refolding) of diluted proteins. Also contribute to study protein solubility ^[15]^.

## Supporting Information

The authors have cited additional references within the Supporting Information ^[12a, 16] **[11, 17] [18] [19] [20] [21] [22] [23]**^

## Acknowledgements

Department of Dermatology and Venerology, Aalborg University Hospital, Hobrovej 18-20, Aalborg 9000, Denmark.

## Supporting Information

### Stabilizing role of HSA

BOCOUTURE preparation has HSA as stabilizer molecule and botulinum toxin as active principle. The stabilizers can be a molecule of low molecular weight such as caprylate and acetyltryptophanate in the case of commercially available HSA ^[1]^ or a highly stable protein with a well-defined structure such as HSA ^[2, 3]^ which has been used to stabilize proteins in lyophilized formulations ^[4]^ (case of BOCOUTURE). The mechanism of action of these stabilizer molecules follows different rules: they interact directly with the active substance (caprylate and acetyltryptophanate) ^[1]^ or the active principle can be shielded to avoid self-aggregation, as HSA does with BoNT-A ^[5]^, in this case, its concentration is higher than the active principle.

The stabilizing role of HSA is based on its high stability. Thus, HSA can maintain the structural integrity of the active principles (BoNT-A) acting as a scaffold to prevent conformational changes ^[6, 7]^. In addition, reduces the likelihood of aggregation triggered by self-interactions ^[5]^ and stabilize the pH within the formulation ^[8, 9]^.

The BoNT-A is clinically utilized for diverse conditions including chronic migraine, cervical dystonia, muscle spasticity, strabismus, overactive bladder, and hyperhidrosis ^[10, 11]^. In aesthetic dermatology, it is extensively employed for reducing facial wrinkles, such as frown lines, crow’s feet, and forehead lines, as well as for softening neck wrinkles and marionette lines ^[12, 13]^. Thus, formulation improvements are a central issue in the pharmaceutical industry aimed at clinical applications. Two crucial points are highlighted in the formulation field: 1) the stabilizer role in preserving the activity of the active substance, and 2) the sensitivity of the methodology employed to conduct stability and quality control studies.

### Single exponential decay to evaluate the time constant

Passing the melting point, the behavior following a single exponential decay kinetic (**τ**_**1pM**_ **= 67**,**07 sec and τ**_**1nM**_ **= 529 sec**) according to MATLAB calculations, the general model for 1pM is giving by the next equation: **f(x) = a+b*exp(c*(x-1200)**, where **a** = -4.429×104, **b** = 8445, **c** = -0.01491 with a 95% of confidence and **r**^**2**^ = 0.9771 (**Fig. 1C**), 1200 denotes the location of the peak of the thermal transition.

## Experimental Details

### Materials

Bovine serum albumin (BSA) (Lot: SLBM5216V) was acquired from Sigma-Aldrich (St. Louis, MO, USA). Louis, MO, and was dissolved in both phosphate-buffered saline (PBS) at pH7 from VWR, BDH Chemicals (Lot:23F214144) or water Milli Q Type II was used. Quartz cuvettes acquired from Hellma Analytics (need an extreme cleaning process in order to work at ultralow proteins concentration) and Acrylic cuvettes from Sarstedt (Germany). Human Serum Albumin (HSA) (Lot: P100491951) was acquired from CSL Behring GmbH, Tyskland, Germany, and Dry Botulinum Toxin formulation (BOCOUTURE) (Lot: 139059) from Merz Pharmaceuticals GmbH, 60318 Frankfurt/Main Tyskland, Germany; both preparation was dissolved in NaCl 0.9% (Lot: 22452413) from B. Braun Melsungen AG 34209 Melsungen, Germany.

### Fluorescence spectra

Fluorescence spectra were recorded using a Horiba Scientific Spectrofluorometer Model with a 75 W Xenon ARC lamp (Ushio Inc. Japan) in a cuvette holder with a temperature control system at 25 °C, and a 10 × 2 mm quartz cell designed for a low volume. The excitation and emission wavelengths were 295 nm and 350 nm, respectively, with excitation slit of 1,4 mm and 10 nm, and emission slits of 20 mm and 30 mm, respectively, to improve the light input in the PMT. The integration time was 10 s and the scan rate was 1 nm/s.

### Determination of the HSA Concentration in the BOCOUTURE Sample and preparation of BT from dry formulation

Dry botulinum toxin from Merz Pharmaceuticals GmbH (BOCOUTURE®) was prepared in NaCl 0.9%. According to the International Unit System, 20 IU = 1 ng. One vial containing 50 IU (2.5 ng) of BT was dissolved in 0.005 L of 0.9% NaCl to obtain 3.33 pM of BoNT-A. To determine the concentration of Human Serum Albumin (HSA) in the BOCOUTURE sample, a calibration curve method was employed. The BOCOUTURE sample contained both HSA (unknown concentration) and BoNT-A at 3.3 pM, allowing the HSA concentration to be calculated by subtracting the BoNT-A concentration from the total protein concentration.

### Preparation of HSA Calibration Curve

Pure HSA (Human Serum Albumin) was used to create a calibration curve (r^2^ = 0.99) for measuring absorbance at 280 nm. Different dilutions of pure HSA were prepared using a NaCl 0.9% solution. These dilutions were prepared meticulously to ensure accurate measurements. The curve allowed the calculation of HSA concentration [HSA] = [Total Protein]-[BONT-A] = 2.8 nM in the BOCOUTURE formulation, which is essential for the accurate determination of the melting point of the HSA stabilizer.

### Absorbance Measurements

Absorbance measurements of the HSA dilutions were carried out at 280 nm using a precision absorbance instrument DS5 UV-VIS spectrophotometer from Edinburgh Instruments Ltd, United Kingdom. The instrument was calibrated according to the manufacturer’s specifications. In addition, before the measurements were taken, a baseline was made.

### Calculation of HSA Concentration

The absorbance values obtained from the HSA dilutions were used to generate a standard calibration curve by plotting the absorbance versus known HSA concentrations. This curve served as a reference to determine the HSA concentration in the BOCOUTURE sample (**see Figure 1**).

**Figure S1:**
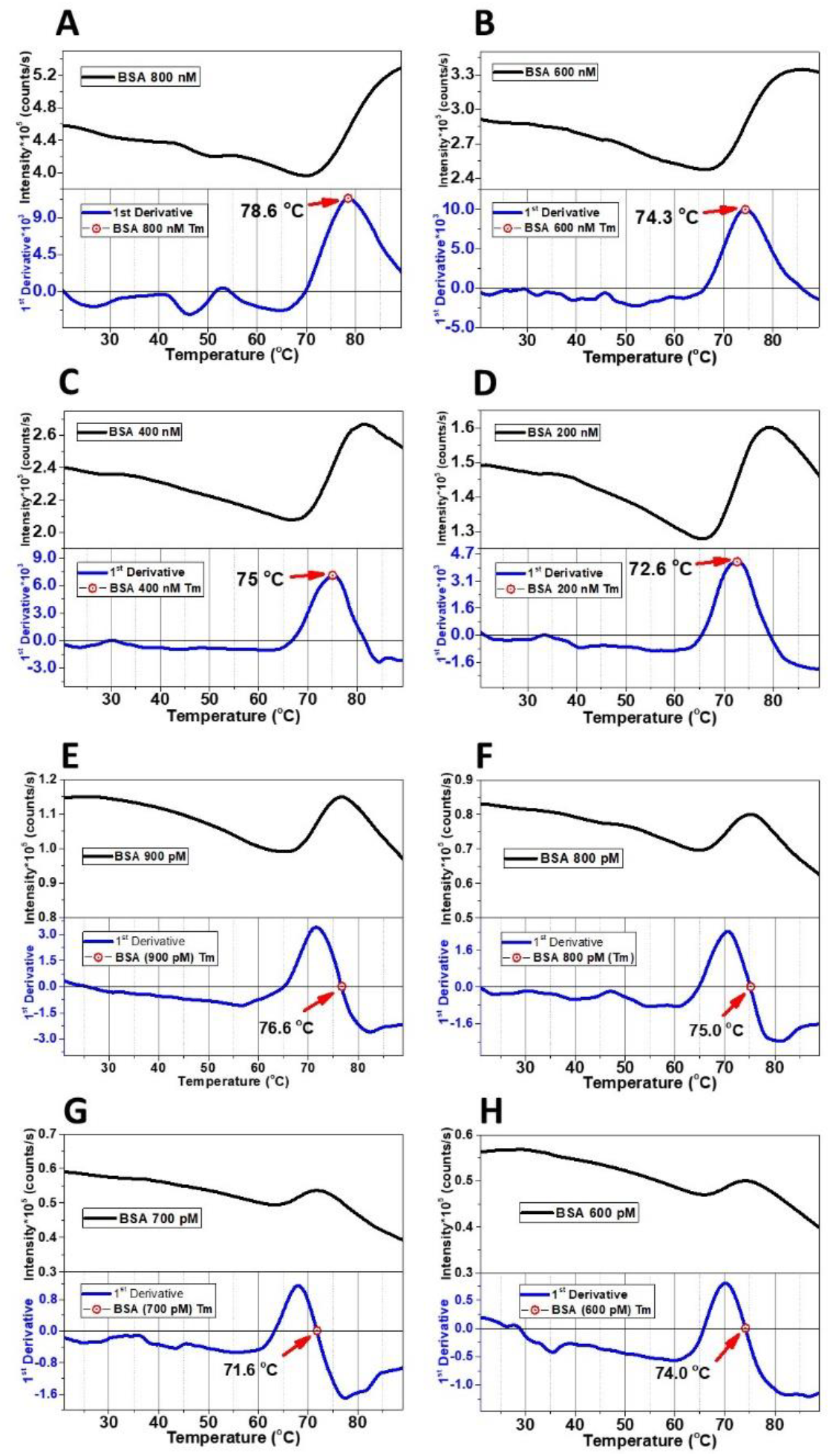
Resonance light scattering to detect the melting point at additional concentration. From 200 nM to 800 nM (A-D) and from 900 pM to 600 pM (E-H).

### Measurement of Total Protein Concentration in BOCOUTURE Sample

A sample of the BOCOUTURE dry formulation, containing both HSA (unknown concentration) and BoNT-A at 3.3 pM, was subjected to absorbance measurements at 280 nm. The measured absorbance corresponded to the total protein concentration of the sample, which included both HSA and BoNT-A.

### Calculation of HSA Concentration in BOCOUTURE

Given that the concentration of BoNT-A in the BOCOUTURE sample was known to be 3.3 pM, the HSA concentration in the BOCOUTURE sample was determined by subtracting the BT concentration from the total protein concentration measured, which was 2.8 nM. This simple calculation provides the HSA concentration for pharmaceutical formulations.

This simple method allowed for accurate quantification of HSA in the BOCOUTURE sample, facilitating the subsequent analysis of protein stability and melting point determination in the context of this pharmaceutical product.

### Melting point determination

The melting point of both BSA and BoNT-A were recorded in the same instrument to obtain the fluorescence spectra using a scattering setting couplet to a ramp temperature. The temperature was varied from 20 °C to 75/80 °C in pure milli Q water and PBS pH7. The excitation emission wavelength was 295 nm. The S/N of the spectra were improved by using 0,1 points/sec in the recording. The accuracy of the melting point is shown by the first derivative.

Removing bubbles in the sample: The reduction in the bubble content in the sample was enhanced by 1- the Vacuum degassing process, which minimized potential sources of interference in the scattering analysis.

2- Heating process in the same temperature range. Increasing the temperature reduced the solubility of the gases in the solution, facilitating the escape of bubbles from the solution to the air ^[14]^.

